# A reference-quality, fully annotated genome from a Puerto Rican individual

**DOI:** 10.1101/2021.06.10.447952

**Authors:** Aleksey Zimin, Alaina Shumate, Ida Shinder, Jakob Heinz, Daniela Puiu, Mihaela Pertea, Steven L. Salzberg

## Abstract

Until 2019, the human genome was available in only one fully-annotated version, GRCh38, which was the result of 18 years of continuous improvement and revision. Despite dramatic improvements in sequencing technology, no other genome was available as an annotated reference until 2019, when the genome of an Ashkenazi individual, Ash1, was released. In this study, we describe the assembly and annotation of a second individual genome, from a Puerto Rican individual whose DNA was collected as part of the Human Pangenome project. The new genome, called PR1, is the first true reference genome created from an individual of African descent. Due to recent improvements in both sequencing and assembly technology, and particularly to the use of the recently completed CHM13 human genome as a guide to assembly, PR1 is more complete and more contiguous than either GRCh38 or Ash1. Annotation revealed 37,755 genes (of which 19,999 are protein-coding), including 12 additional gene copies that are present in PR1 and missing from CHM13. 57 genes have fewer copies in PR1 than in CHM13, 9 map only partially, and 3 genes (all non-coding) from CHM13 are entirely missing from PR1.

## Introduction

Since 2001, most work in human genomics and genetics has relied upon a single genome, which was originally sequenced from a small number of anonymous individuals of varying, unknown backgrounds [1]. The human reference genome, currently in release GRCh38, is a mosaic of those individuals, originally created by stitching together individual assemblies of thousands of short bacterial artificial chromosomes (BACs), which averaged ∼150 kilobases (Kb) in length.

The human reference genome is the basis for an enormous range of scientific and medical studies, and for large databases of genome variation such as gnomAD [2]and dbSNP [3] that aim to catalog the genetic variation inherent in diverse populations. Although thousands of human genomes have been sequenced in the past decade, until recently none of them were available to function as reference genomes, for at least two reasons. First, none were as contiguous as GRCh38, which has undergone many years of careful refinement to fill gaps and fix errors. This is now changing as a result of extremely long reads that can be generated by the latest sequencing technology, allowing us to generate assemblies that are more complete than GRCh38. Second, an essential requirement for a reference genome is that it is annotated with all known human genes, including both protein-coding and noncoding genes. Until the publication of the Ash1 genome in 2019 [4], no human genome other than GRCh38 was both highly contiguous and fully annotated.

A distinctive feature of Ash1 is that it is based on a single individual of Ashkenazi descent, unlike GRCh38 which was created from a mixture of multiple individuals with different genetic backgrounds. For some genetic studies, particularly those focused on a specific population, an ideal genetic reference would be a genome assembled from a healthy individual from a similar genetic background. One reason for this, as discussed in [4], is that comparisons between new study subjects and an appropriately matched reference will turn up many fewer variants, allowing investigators to focus on those variants most likely to be relevant. The Ash1 genome demonstrated that it was technically possible to create such reference genomes, and that more are needed.

The current study describes the assembly and annotation of PR1, the first genome of a Puerto Rican individual, and the first reference-quality genome of any human of African descent. As we show in this study, the contiguity and completeness of PR1 are considerably better than GRCh38, with more total DNA, fewer gaps, and more genes. For these and other reasons, PR1 may be preferable to use as a reference genome in studies of individuals from a similar Puerto Rican genetic background.

## Results and discussion

### Data and genetic background

The data for the HG01243 genome was generated and made publicly available by the Human Pangenome project (https://github.com/human-pangenomics/hpgp-data), which in turn used data and samples from the 1000 Genomes Project [5] and the Genome In A Bottle (GIAB) [6] collections. Several earlier assemblies of HG01243 appeared as part of an assembly methods comparison study [7]; we refer to our assembly as PR1. HG01243 was chosen for the current study because the individual is distinct from the only previous individual human reference-quality assembly, that of an Ashkenazi individual [4], and because he represents a population, Puerto Ricans, that is not well-represented by the GRCh38 reference genome. Another benefit of choosing HG01243 is that sequence data is available from both parents, HG01241 and HG01242, which offers future opportunities to study recombination and genetic differences between parents and offspring.

HG01243 is a male individual from Puerto Rico of mixed, primarily African, ancestry. His maternal haplogroup based on the mitochondrial genome is L1b2a, which is most common in West Africa [8]. The paternal haplogroup based on the Y chromosome is R1b1a1b (more specifically, R1b1a1b1a1a), which is the most common haplogroup in western Europe [9]. We compared HG01243’s Y chromosome to males of diverse backgrounds in the 1000 Genomes (1KG) and Human Genome Diversity Project (HGDP) callset available from gnomAD v3.1 [2] and determined that his Y chromosome has the highest similarity to males of Basque descent, which is consistent with the R1b1a1b haplogroup.

To infer PR1’s ancestry from the whole-genome data, we followed an approach developed by Stevens et al. [10] that uses identity-by-descent (IBD) and identity-by-state (IBS) to determine the extent of relatedness between a pair of individuals. Briefly, in a pairwise comparison of two individuals at a single locus or allele, there are three possible IBS outcomes: IBS0, IBS1, and IBS2. Loci where the individuals do not have any alleles in common (for example, a pair of individuals with genotypes AA and BB) are labeled IBS0. Loci where the individuals share a single allele (e.g., AA and AB) are IBS1, and loci where individuals share both alleles (e.g., AA and AA) are IBS2. A subtype of IBS2 called IBS2* [10] occurs when two individuals are heterozygous and share both alleles (i.e., AB and AB). Pairwise relationships for all individuals in a given cohort can be plotted by integrating IBS values over a set of variants. These relationships for PR1 are shown in Figure 1, where the x-axis ratio uses the IBS2* ratio, IBS2*/(IBS2* + IBS0) [10]. For unrelated individuals from the same population, where all loci are assumed to be biallelic and follow Hardy-Weinberg equilibrium, the IBS2* ratio is expected to average ⍰, whereas the IBS1 heterozygosity ratio (the y-axis of Figure 1) is expected to average 1 [11]. As shown in Figure 1, the IBS2* ratio for pairwise comparisons of PR1 to African-descent individuals (shown in yellow) averages just below ⍰, indicating that PR1 is of African ancestry.

**Figure 1.**
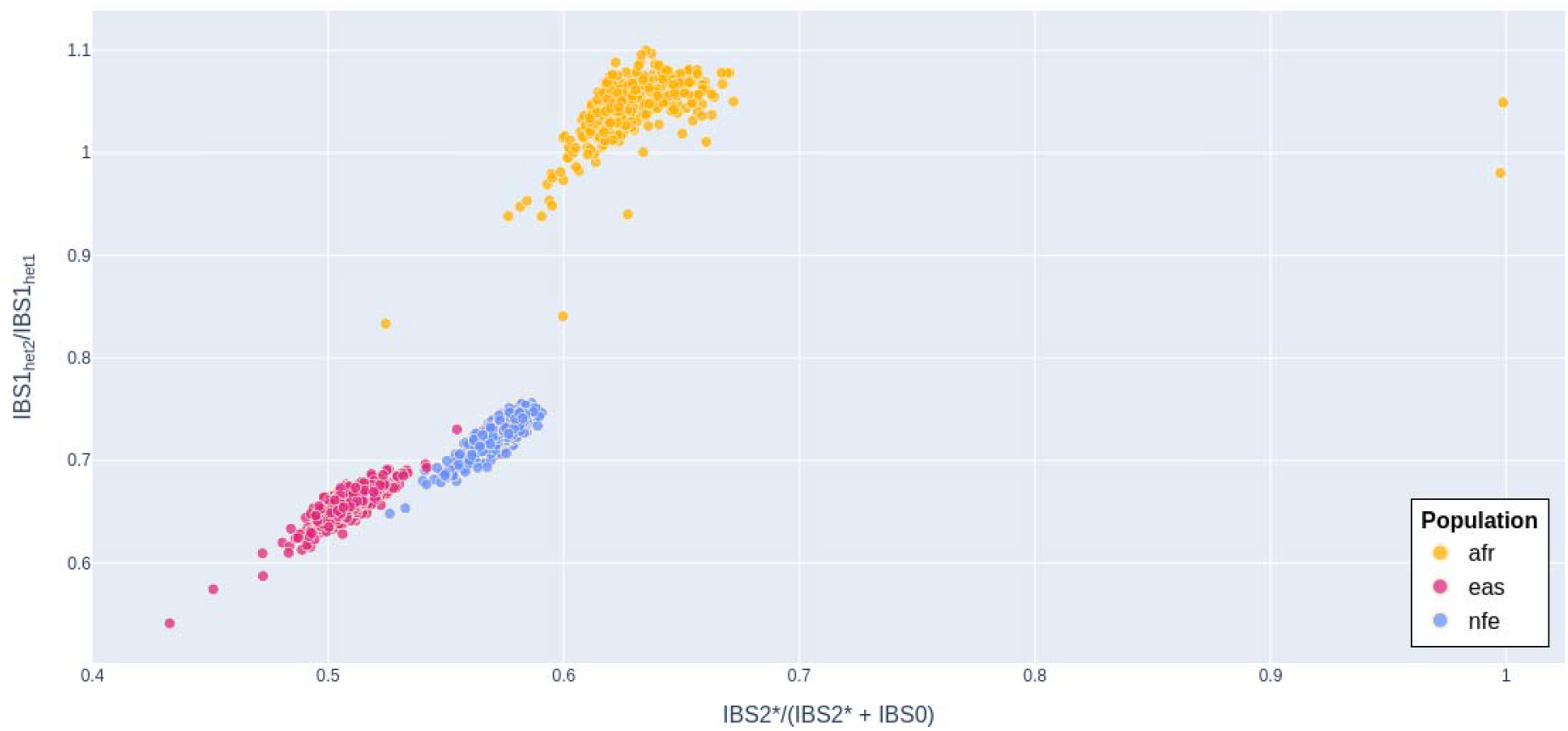
IBS relationship between PR1 and 2,371 individuals of African/African-American (afr), Non-Finish European (nfe), and East Asian (eas) descent in the 1KG and HGDP callset. Population labels were obtained from gnomAD v3.1 [2]. The x-axis ratio has two components, IBS0 and IBS2*. IBS0 is an aggregate count of times that a given individual does not share any alleles with PR1 (i.e., PR1 is AA and the target individual is BB). IBS2* is the aggregate count of times that the target individual is heterozygous and shares two alleles with PR1 (genotypes AB and AB). The y-axis ratio (IBS1_het2_/IBS1_het1_), is a proxy for heterozygosity and level of genetic variation. IBS1_het1_ is the aggregate count of times that PR1 is heterozygous and the target individual is homozygous and they share one allele (i.e., AB and AA, respectively). IBS1_het2_ is the aggregate count of times that PR1 is homozygous and the target individual is heterozygous and they share one allele (i.e., AA and AB, respectively). As expected for parent-child relationships, the IBS2* ratio for PR1’s parents, who appear as two dots on the far right of the plot, is 1.

### Assembly

To create the PR1 assembly, we used both long and short reads from multiple technologies, including PacBio continuous long reads (CLR), PacBio high-fidelity reads (HiFi), Oxford Nanopore (ONT) ultra-long reads, and 150bp Illumina paired-end reads (**Table 1**). Our goal was to produce a high-quality, haplotype-merged assembly with a single sequence representing each chromosome. For long-range scaffolding, we used as a reference the assembly of CHM13 released by the Telomere-to-Telomere (T2T) consortium, a very high-quality human assembly that is far more complete and contiguous than the GRCh38 reference genome, with only 5 gaps in version 1.0 [12] [13] and zero gaps in version 1.1. We used the gap-free v1.1 as a guide for our assembly.

**Table 1.** Data used for assembly of the PR1 genome. N50 read length is defined as the length such that 50% of the total sequence is contained in reads of length N50 or longer. Note that the effective length of the Illumina reads was shorter after removing barcodes and adapter sequences.

| Sequence type | Number of reads | N50 read length (bp) | Genome coverage |
| --- | --- | --- | --- |
| Illumina paired-end | 1,307,137,030 | 150 | 59x |
| PacBio CLR | 23,672,554 | 28,837 | 113x |
| PacBio HIFI | 5,485,642 | 19,528 | 34x |
| ONT Ultralong | 16,293,849 | 44.052 | 35x |

We initially produced three different de novo assemblies using three different assemblers, and evaluated them for contiguity and consensus accuracy. To create these assemblies, we used Hifiasm [14] version 0.13-r308 with PacBio HiFi data, Flye [15] version 2.5 with Oxford Nanopore data, and MaSuRCA [16] version 4.0.1 with Illumina, Oxford Nanopore and PacBio CLR data. (Note that for the MaSuRCA assembly, we only used PacBio CLR reads that were longer than 20Kb. A MaSuRCA assembly that included the HiFi reads contained slightly more errors and was not used.) The exact commands used to create each assembly are listed in the Supplementary text. **Table 2** contains summary statistics for these three assemblies. We measured consensus accuracy by aligning the Illumina data to the assemblies and calling variants using Freebayes [17]. A variant was considered an error if no Illumina reads supported the assembly, and if at least 2 Illumina reads supported an alternative base. The MaSuRCA assembly had the best consensus accuracy, while the Flye assembly had the best contiguity, with an N50 contig size of 27.4 Mbp.

**Table 2.** Statistics for the preliminary PR1 assemblies. N50 values were calculated using an estimated genome size of 3.2Gbp; thus an N50 size of 6.7 Mbp means that 1.6 Gbp of the total assembly was contained in contigs at least that large. ONT: Oxford Nanopore Technology.

|  |  |  |  |  |
| --- | --- | --- | --- | --- |
| Assembler | Data used | Assembled sequence | Contig N50 | Number of contigs |
| Hifiasm | PacBio HiFi | 3,096,276,394 | 6,760,611 | 1,473 |
| MaSuRCA | Illumina+ONT<br>+PacBio CLR >20Kb | 2,848,911,829 | 7,622,810 | 1,496 |
| Flye | ONT | 2,906,307,381 | 27,370,234 | 1,693 |

From these initial assemblies, we chose the Flye assembly for subsequent refinements because it had the best contiguity. To improve base-level accuracy, we merged the MaSuRCA assembly with the Flye assembly to use the MaSuRCA consensus where possible (see Methods). The resulting MaSuRCA+Flye merged assembly contained 2,892,693,836 bp in 1,185 contigs. Note that this assembly merging process has advantages over simply “polishing” one assembly using Illumina data, because the large contigs in the assemblies can be mapped unambiguously in many places where short Illumina reads would not map reliably enough for accurate polishing.

We then used the MaSuRCA chromosome scaffolder [4] to order and orient the merged contigs onto the human reference using chromosomes 1-22 and X from the CHM13 v1.1 reference genome (https://github.com/nanopore-wgs-consortium/CHM13) produced by the T2T consortium [12] [18], and using the Y chromosome from HG002 that was recently finished by the same consortium. This is similar to the procedure used to build the Ashkenazi reference genome, with the major difference being that Ash1 used GRCh38 as the primary backbone for scaffolding. (See Methods for more details of the scaffolding process.) The final reference-guided assembly has each chromosome in a single scaffold and contains only 64 unplaced contigs containing 1.09 Mbp of sequence (**Table 3**).

**Table 3.**
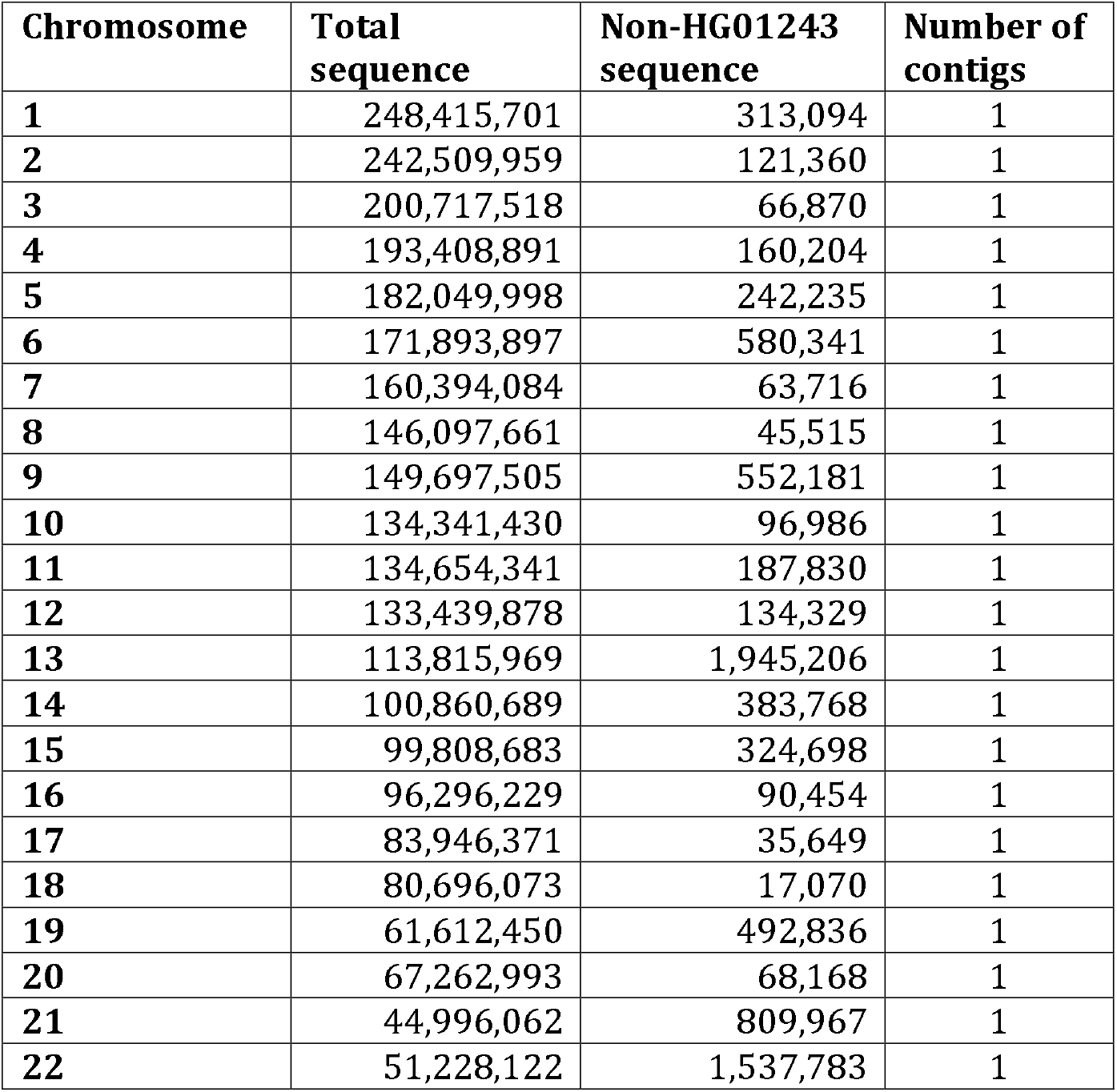
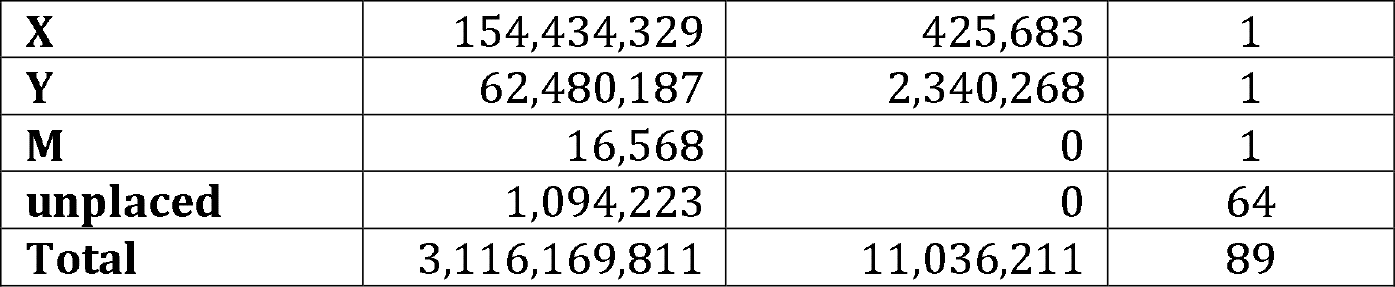
Chromosome sizes and the amount of non-HG01243 sequence per chromosome in the final PR1 assembly.

|  |  |  |  |
| --- | --- | --- | --- |
| **Chromosome** | **Total sequence** | **Non-HG01243 sequence** | **Number of contigs** |
| **1** | 248,415,701 | 313,094 | 1 |
| **2** | 242,509,959 | 121,360 | 1 |
| **3** | 200,717,518 | 66,870 | 1 |
| **4** | 193,408,891 | 160,204 | 1 |
| **5** | 182,049,998 | 242,235 | 1 |
| **6** | 171,893,897 | 580,341 | 1 |
| **7** | 160,394,084 | 63,716 | 1 |
| **8** | 146,097,661 | 45,515 | 1 |
| **9** | 149,697,505 | 552,181 | 1 |
| **10** | 134,341,430 | 96,986 | 1 |
| **11** | 134,654,341 | 187,830 | 1 |
| **12** | 133,439,878 | 134,329 | 1 |
| **13** | 113,815,969 | 1,945,206 | 1 |
| **14** | 100,860,689 | 383,768 | 1 |
| **15** | 99,808,683 | 324,698 | 1 |
| **16** | 96,296,229 | 90,454 | 1 |
| **17** | 83,946,371 | 35,649 | 1 |
| **18** | 80,696,073 | 17,070 | 1 |
| **19** | 61,612,450 | 492,836 | 1 |
| **20** | 67,262,993 | 68,168 | 1 |
| **21** | 44,996,062 | 809,967 | 1 |
| **22** | 51,228,122 | 1,537,783 | 1 |
| **X** | 154,434,329 | 425,683 | 1 |
| **Y** | 62,480,187 | 2,340,268 | 1 |
| **M** | 16,568 | 0 | 1 |
| **unplaced** | 1,094,223 | 0 | 64 |
| **Total** | 3,116,169,811 | 11,036,211 | 89 |

We closed gaps in the chromosome scaffolds by aligning the MaSuRCA contigs to the scaffolds with nucmer, retaining the unique 1-to-1 alignments from delta-filter [19], and closing each gap where the MaSuRCA contig was anchored by unique alignments of 5 Kbp or longer on both sides of the gap. This step closed 147 gaps. We filled in the remaining gaps in the chromosome scaffolds using the CHM13 genome sequence, which we recorded in lowercase while HG01243 sequence was uppercase. Essentially all gaps that were filled in this way represented hard-to-assemble repetitive sequences. We included a total of 280 Mbp of sequence from CHM13 and HG002 (for chrY only) in the final 3.1 Gbp assembly. We followed the gap filling by two rounds of polishing with Nextpolish [20], using PacBio HiFi data for the first round and Illumina data for the second. Most of the sequences used to fill gaps were confirmed by PR1 reads during these polishing steps, and those bases were converted to uppercase.

After the final polishing step, the assembly had 3,115,075,588 bp of sequence on the 24 chromosomes and the mitochondrial genome, plus 1,094,223 bp in 64 unplaced contigs, for a total of 3,116,169,811 bp. In the final PR1 assembly, all chromosomes are complete, assembled end-to-end with no gaps. All chromosome sizes and the amount of non-PR1 sequence are listed in **Table 3**. The Non-HG01243 column shows the number of bases from the T2T CHM13 reference in each chromosome that we were unable to replace with read alignments from the HG01243 data.

### Validation

We performed validation checks on the assembly using Merqury [21] with the combined set of Illumina and PacBio HiFi data. Merqury uses the reads to identify all “good” k-mers (here we used k=25), and then marks any k-mer in the assembly that does not appear in the reads as bad. After masking out the CHM13-derived sequences, we used Merqury to calculate a base-level quality score (QV) of 70.1, which corresponds to an error rate of just under 1 error per 10 million bases.

For additional validation we mapped the entire PacBio HiFi dataset to the assembly using minimap2. We then computed the mapped coverage of the HiFi reads at every base and plotted that as a histogram, shown in Supplementary **Figure S1**. The main peak is at coverage ∼37, which corresponds to the expected coverage by the HiFi reads. There is a small second peak at 18, which corresponds to regions where the two haplotypes diverge, and where we expect to see one-half of the total coverage.

As an independent validation of the scaffold structure of PR1, we used chromosome conformation capture (Hi-C) data from HG01243 to build a contact map, shown in **Figure S2**. As shown in the map, the Hi-C data is in close agreement with the overall genome structure for all chromosomes, with most contacts clustering along the main diagonal as expected for a correct assembly.

### Novel sequence in PR1

In addition to the sequences placed on chromosomes, the PR1 assembly has 64 small contigs containing 1,094,223 bp that could not be placed and that are non-repetitive, representing sequences that are unique to PR1. 42 of these contigs totaling 297 Kb do not align elsewhere in the genome (defined as not having any alignment that covers more than 5% of the contig), while the remaining 18 contigs align partially, ranging from 5–75% of their length.

To identify additional novel sequence in PR1, we aligned all chromosomes in the assembly to the corresponding CHM13 chromosomes. After removing simple sequence repeats and other recognizably repetitive sequences, we were left with 14.5 Mbp in 5,047 insertions in PR1, with lengths ranging from 200–203,707 bp (of which 10 insertions were longer than 100 Kb). When we add in the unplaced contigs, the PR1 genome contains ∼15.6 Mb of novel sequence, likely representing regions that are distinctive in the Puerto Rican population.

### Comparison to other reference genomes

PR1 is the third fully-annotated reference genome to be created, and we compared it to the longstanding GRCh38 reference as well as to the Ashkenazi reference genome, Ash1. Because the Ash1 assembly used GRCh38 as a template, both it and GRCh38 are less complete than PR1, which used the complete CHM13 assembly as a guide. For our comparisons here, we only considered regions that are present in all three assemblies, and we excluded any sequence in PR1 that was filled in using CHM13.

Tables 4 and 5 show a summary of these findings. In Table 4, which compares PR1 and Ash1 to GRCh38, we see that PR1 has a total of 6,550,479 variants as compared to GRCh38, out of which 3,512,566 are heterozygous variants where one of the haplotypes agrees with GRCh38. In contrast, the Ashkenazi genome has fewer differences, 5,984,422 in total, of which 2,719,200 are heterozygous sites where one variant agrees with GRCh38.

**Table 4.** PR1 and Ash1 variants compared to GRCh38. Variants include both substitutions and indels (up to 200bp) in Ash1 or PR1 compared to GRCh38. We used the original Illumina reads to determine whether variants in PR1 and Ash1 were homozygous or heterozygous.

Tables 4 and 5 show a summary of these findings.
|  |  |  |
| --- | --- | --- |
| Variant type | Both variants disagree with GRCh38 | One variant agrees with GRCh38 |
| PR1 homozygous | 3,021,169 | n/a |
| PR1 heterozygous | 16,744 | 3,512,566 |
| Ash1 homozygous | 3,249,658 | n/a |
| Ash1 heterozygous | 15,564 | 2,719,200 |

**Table 5.** Comparison between Ash1 and PR1 at sites where both differ from GRCh38 and where they differ from each other. Most of the locations correspond to variants that are homozygous in both genomes. For cases where both genomes are homozygous or both are heterozygous, numbers in parentheses indicate the number of positions where PR1 and Ash1 match each other.

|  | Ash1 homozygous | Ash1 heterozygous |
| --- | --- | --- |
| PR1 homozygous | 1,278,122 (1,220,747) | 393,797 |
| PR1 heterozygous | 551,925 | 608,115 (595,395) |

In Table 5, we compare PR1 and Ash1 directly at 2,223,844 locations where they both differ from GRCh38. In 1,278,122 of these locations, both PR1 and Ash1 are homozygous, and at 1,220,747 (96%) of these sites, PR1 and Ash1 agree with each other, suggesting that GRCh38 has a less-common variant at those sites. Similarly, the two genomes agree on both variants at 593,395 of the 608,115 sites (98%) where both PR1 and Ash1 are heterozygous but disagree with GRCh38. Data from the heterozygous sites also suggest that PR1 is more heterozygous than Ash1, because it has a greater number of heterozygous sites than Ash1 when they both have a variant compared to GRCh38 at the same location.

PR1 should serve well as a reference genome in studies of other Puerto Rican individuals of similar genetic ancestry. As a proof-of-concept experiment to illustrate this point, we computed variants including SNPs and small indels from HG01241 and HG01242, who are the parents of HG01243, using first PR1 and then GRCh38 as the reference. As shown in Table 6, we found approximately half as many variants in either the father (52%) or the mother (48%) when using PR1 as the reference compared to GRCh38. In general, individuals who are closer genetically to PR1 than to other available references will have considerably fewer variants, and this in turn may narrow the search for the causes of genetic disorders in those individuals.

**Table 6.** Number of variants found when comparing the parents of PR1, HG01241 and HG01242, to the PR1 and GRCh38 genomes.

|  | Substitutions | Insertions/Deletions | Total |
| --- | --- | --- | --- |
| PR1 vs. HG01241 | 786,619 | 688,552 | 1,475,171 |
| PR1 vs. HG01242 | 691,058 | 607,832 | 1,298,890 |
| GRCh38 vs. HG01241 | 1,430,509 | 1,426,556 | 2,857,065 |
| GRCh38 vs. HG01242 | 1,356,876 | 1,342,086 | 2,698,962 |

### Gene Annotation

We used Liftoff [22] to map all of the genes from CHM13 onto PR1, including protein-coding and non-coding RNA genes. The CHM13 annotation contains 37,670 genes in total, of which 19,829 protein-coding genes and 16,818 lncRNAs (36,647 genes) were mapped onto CHM13 from the GENCODE annotation of GRCh38 [23]. CHM13 also contains 804 additional paralogs (140 protein-coding and 664 lncRNAs) and 219 additional rDNA genes not present in GRCh38. Because the CHM13 genome does not have a Y chromosome, we mapped the GENCODE genes from GRCh38’s chromosome Y onto PR1 chromosome Y.

Out of the 37,670 genes from CHM13 and 142 genes from GRCh38 chrY (37,812 total), Liftoff successfully mapped 37,743 (99.8%). Of the 69 unmapped genes (shown in Supplementary Table S1), 42 are protein-coding and 27 are non-coding. Liftoff considers a gene to be mapped if the alignment coverage and sequence identity of the exons are greater than or equal to 50%; however, the vast majority of genes in PR1 greatly exceed this threshold, with 93% of genes mapping with >= 99% coverage and sequence identity.

Out of the 69 genes that failed to map, 29 aligned end-to-end with another copy of the gene present elsewhere in the assembly (i.e., a paralog), suggesting that PR1 simply has fewer copies (Table S1). Another 28 genes had partial copies present in the assembly (see Methods). Of the 12 remaining unmapped genes, all but 3 genes mapped partially but did not meet the 50% minimum coverage and sequence identity threshold. The 3 genes completely missing from PR1 are all lncRNAs whose function is unknown.

We looked at all 86,335 protein-coding transcripts that were mapped from CHM13 to PR1 to determine if the protein sequence was preserved. In the vast majority of cases, the sequences were either identical or had non-synonymous mutations that preserved the protein sequence length. Specifically, 71,699 transcripts (83.0%) had identical sequence, 13,544 (15.7%) had amino acid changes but identical lengths, and 828 (0.96%) had insertions or deletions that preserved the reading frame. Only 196 transcripts had frame-shifting mutations, and 68 were truncated on one end or missing the start codon.

### Extra gene copies in PR1

To identify genes with a higher copy number in PR1 than CHM13, we used an optional feature of Liftoff to identify additional paralogs. We found 12 additional paralogs including 8 paralogs of protein-coding genes and 4 paralogs of lncRNAs (Table S2). Six of these paralogs occur in tandem, defined as a gene that occurs within 100 Kbp of another copy. All isoforms of the additional copies are 100% identical at the mRNA level to the original copy in CHM13. In general, a finding of additional paralogs is either the result of increased assembly completeness or copy number variation. Given that CHM13 is a complete, gap-free assembly, these 12 paralogs appear to represent genuine copy number variation between PR1 and CHM13. Also worth noting here is that CHM13 contains 140 additional copies of protein-coding genes by comparison to GRCh38 [18], all of which are also present in PR1.

Because GRCh38 is currently the primary human reference genome, we also mapped the annotation from GRCh38 onto PR1, using CHESS v2.2. The CHESS annotation [24] includes all protein-coding genes from both GENCODE [23] and RefSeq [25], but the noncoding genes are substantially different, as are the precise splice variants for many protein-coding genes. We successfully mapped 42,172 out of 42,306 genes (99.7%) from the CHESS annotation. 73 out of the 134 unmapped genes are protein-coding and the other 61 are non-coding. We also identified 159 additional gene copies (paralogs) present in PR1 and missing from GRCh38. These include 30 paralogs of protein-coding genes and 129 paralogs of non-coding genes. The CHESS genes that failed to map, including all gene types, are shown in Supplementary Table S3. All extra gene copies in PR1 as compared to both CHM13 and GRCh38, along with the gene names and chromosomal locations on PR1, are shown in Supplemental Tables S2 and S4.

## Methods

### Assembly polishing and scaffolding

To improve base-level accuracy, we aligned the MaSuRCA assembly to the Flye assembly using Nucmer [26], and identified the alignments with 1-to-1 best matches using delta-filter. We merged alignments when a MaSuRCA contig matched a Flye contig in several chunks that had the same order and orientation, producing spanning alignments that started at the beginning of the first chunk and ended at the end of the last chunk. Next, we replaced the Flye assembly’s consensus sequence with the higher-quality MaSuRCA consensus in places where MaSuRCA and Flye contigs aligned. This process replaced 99.4% of the Flye sequence with the MaSuRCA consensus.

Next, we added all MaSuRCA contigs that aligned to the Flye assembly over less than 25% of their length, on the assumption that these contigs contained sequence missing from the Flye assembly. This process added 90 MaSuRCA contigs containing 2,263,294 bp to the assembly. We aligned the assembly to itself and removed any contigs that mapped over 90% of their length to the interior of larger contigs. These contigs were likely to represent redundant haplotype copies; i.e., places where the two haplotypes diverged sufficiently that the assembly algorithm created two separate contigs.

To scaffold the contigs, we used the chromosome scaffolder pipeline distributed with MaSuRCA assembler. The chromosome scaffolder first aligned the assembly to the CHM13 reference genome and automatically identified breaks in the alignments where the mapping indicated a possible mis-assembly. Each such location was then checked by mapping the underlying reads to the assembly and examining read coverage. If read coverage dropped below 10% of the average coverage C within 50Kb of the breakpoint, the assembled contig was assumed to be erroneous and was broken at the lowest coverage point. If read coverage was too high (>C/ln(2)), suggesting a repeat-induced mis-assembly, we looked for two breakpoints flanking the high-coverage region, and split the assembly at both breakpoints.

This process introduced 703 gaps (breakpoints) into the MaSuRCA+Flye merged assembly, resulting in 1888 contigs. We were able to close 134 of these gaps after the contigs were placed on the chromosomes, which confirmed that they were correctly assembled. Then we used the CHM13 1.1 assembly (plus chromosome Y from HG002) as a reference to order and orient the contigs, placing each contig where it aligned best. This procedure ordered and oriented 1,359 contigs, containing 98.8% of the sequence, onto chromosome-scale scaffolds. 34.2 Mbp of sequence remained in unplaced contigs. When we aligned these contigs to the CHM13 assembly, we found that 33.0 Mbp of these sequences were collapsed repeat copies that aligned with 90-95% identity to centromeric regions. Because of the high error rates in the Nanopore data, the assembler likely collapsed these repeats producing spurious “consensus” of multiple slightly divergent repeat copies. (Note that these sequences contained no genes.) Thus we decided to exclude these sequences from the assembly.

The last step in assembly was polishing with Nextpolish [20]. The polishing was applied to the entire genome, including the sequences that were filled in using CHM13. We polished the genome in two iterations, first using PacBio HiFi data and then using Illumina data. During this process, many small variants within the CHM13-derived regions where the HG01243/PR1 data aligned unambiguously were corrected to represent the PR1 genome sequence. All bases that were corrected or confirmed by the read alignments are indicated in uppercase in the final assembly, even those within CHM13-derived regions.

To build the contact map from the Hi-C data from HG01243, we used Juicer [27] version 1.6 (https://github.com/aidenlab/juicer/releases) to build the .hic file and visualized it with Juicebox for Windows version 1.11.08 [Durand NC, Robinson JT, Shamim MS, Machol I, Mesirov JP, Lander ES, Aiden EL. Juicebox provides a visualization system for Hi-C contact maps with unlimited zoom. Cell systems. 2016 Jul 27;3(1):99-101.].

### Genotyping

To identify the paternal haplogroup of PR1, we used Yleaf [28] applied to the Y chromosome. To identify the maternal haplogroup, we used Haplogrep 2 [29] on the mitochondrial genome. To compare the PR1, GRCh38, and Ash1 assemblies, we aligned all pairs of genomes using nucmer [26] and then computed SNPs and indels using nucmer’s delta2vcf utility. We then compared these variants to the original Illumina reads for both PR1 and Ash1 in order to identify which variants were homozygous and heterozygous.

To identify variants in PR1’s parents, HG01241 and HG01242, we aligned all Illumina reads from each of those genomes (∼532M reads for HG01241 and ∼574M reads for HG01242) to both PR1 and GRCh38 using bwa-mem [30], and then called variants with FreeBayes [17].

For IBS analysis, we used Hail v0.2.67 [31] to parse the 1KG and HGDP variant files from gnomAD v3.1 (https://cloud.google.com/life-sciences/docs/resources/public-datasets/gnomad). To select high quality variants, we downsampled the 1KG Phase I high-confidence SNPs to randomly keep 1% of the variants, resulting in 281,308 biallelic variants. For each variant, we performed a pairwise comparison of HG01243 alleles to all 3,941 individuals in the 1KG and HGDP call set, with an average of 280,000 non-missing variants per pairwise comparison. Counts of IBS2*, IBS1 and IBS0 for each pairwise comparison were aggregated to calculate the IBS2* ratio (IBS2*/(IBS2* + IBS0) and the IBS1 heterozygosity ratio (IBS1_het2_/IBS1_het1_). To determine the population closest to PR1’s Y chromosome lineage, we filtered the 1KG and HGDP callset for SNPs in the male-specific region of the Y chromosome with a quality value ≥ 20 and calculated the proportion of shared variants to total variants.

### Gene annotation

To annotate the PR1 genome, we mapped the CHM13, GRCh38 chromosome Y, and CHESS v2.2 annotations using Liftoff version 1.6.1 with the following parameters: -copies -polish -exclude_partial -chroms <chroms.txt>. After the initial mapping, we aligned every unmapped transcript to every mapped transcript using Blastn [32] in order to determine if the unmapped genes were copies of mapped genes (where we define a copy as an end-to-end alignment with a mapped transcript). We also re-ran Liftoff allowing for overlapping genes (-overlap 1.0). By comparing the results to the initial Liftoff output, we were able to identify genes that only mapped when allowed to overlap another gene. These overlapping genes are either complete copies of one another if they map to exactly the same locus, or partial copies if they map to different but overlapping loci. The overlapping genes were identified by intersecting the Liftoff-generated annotation file with itself using Bedtools intersect [33]. The output file was then filtered to remove self-overlaps, and genes identified by this process were classified as partial copies. We further attempted to identify partial copies at the protein level by using gffread [34] to extract the protein sequences and blastp [32] to align them to mapped proteins, using an e-value threshold of 10^−6^ to filter the results.

## Data availability

Sequence data including raw signal files and basecalls for HG01243 were previously released by Shafin et al. (20) and are available as an AWS Open Data set for download from https://github.com/human-pangenomics/hpgp-data. Nanopore sequence data are additionally archived at the European Nucleotide Archive under accession code PRJEB37264. The PR1 genome assembly and annotation are available from NCBI and GenBank under BioProject PRJNA730525 and accession GCA_018873775.2, and on GitHub at https://github.com/PuertoRicanGenome. A VCF file listing all of the heterozygous variants in PR1 is available at https://figshare.com/articles/dataset/PR1_v3_0_heterozygous_sites_vcf_gz/16602335.

## Acknowledgements

Thanks to Adam Phillippy, Sergey Koren, and Karen Miga for help with sample selection and data access. This work was supported in part by NIH grants R01 HG006677 and R01 MH123567, and by NSF grants ISO-1744309 and DBI-1759518.

## Notes

https://github.com/PuertoRicanGenome

### Competing Interest Statement

The authors have declared no competing interest.

### Summary of Updates

We used the latest version of chm13 assembly, and the Y chromosomes from the HG002 individual to re-scaffolds the PR1 genome to obtain fully gapless reference. We updated our annotation and report on the fully updated assembly and annotation. We added extra validation steps including analysis by Merqury, and independent validation by HiC data.

