## Supplementary materials for "A reference-quality, fully annotated genome from a Puerto Rican individual"

**Supplementary text and figures**

**Command lines and configurations for creating the initial assemblies of HG01243 (PR1)**.

For the HiFiAsm assembly, we used the following command:

hifiasm -o HG01243.asm -t 32 m64136_200827_191603.fq m64136_200829_012933.fq m64136_200830_075556.fq

The assembly was then converted to fasta using gfatools.

For the Flye assembly, we use the following command:

flye --nano-raw ../Data/nanopore.fa -g 3200000000 -o asm -t 32

For the MaSuRCA assembly, we created the following configuration file:

DATA

PE= pe 500 50 H_IJ-HG01243-HMW-Tube_2-lib1_S2_L001_R1_001.fastq.gz H_IJ-HG01243-HMW-Tube_2-lib1_S2_L001_R2_001.fastq.gz

NANOPORE=nanoporePacbio.20kbmin.fa

END

PARAMETERS

EXTEND_JUMP_READS=0

GRAPH_KMER_SIZE = auto

USE_LINKING_MATES = 0

USE_GRID=1

GRID_ENGINE=SGE

GRID_QUEUE=bigmem.q

GRID_BATCH_SIZE=5000000000

LHE_COVERAGE=25

LIMIT_JUMP_COVERAGE = 300

CA_PARAMETERS = cgwErrorRate=0.15

CLOSE_GAPS=1

NUM_THREADS = 64

JF_SIZE = 200000000

SOAP_ASSEMBLY=0

FLYE_ASSEMBLY=1

END

|  |  |
| --- | --- |
| 0 | 131009333 |
| 1 | 13907632 |
| 2 | 7681203 |
| 3 | 6379153 |
| 4 | 6038061 |
| 5 | 5542162 |
| 6 | 5443038 |
| 7 | 5767905 |
| 8 | 5884768 |
| 9 | 6526640 |
| 10 | 7882978 |
| 11 | 9322434 |
| 12 | 11413723 |
| 13 | 13405819 |
| 14 | 16094044 |
| 15 | 18043050 |
| 16 | 19651441 |
| 17 | 20858732 |
| 18 | 21732866 |
| 19 | 22514578 |
| 20 | 23469886 |
| 21 | 24750872 |
| 22 | 26584901 |
| 23 | 30329151 |
| 24 | 35578741 |
| 25 | 42294203 |
| 26 | 51117986 |
| 27 | 61921968 |
| 28 | 74937100 |
| 29 | 88598442 |
| 30 | 102638633 |
| 31 | 117069418 |
| 32 | 130258439 |
| 33 | 141439906 |
| 34 | 150747708 |
| 35 | 155630615 |
| 36 | 157393676 |
| 37 | 156108699 |
| 38 | 151879425 |
| 39 | 145014305 |
| 40 | 135619249 |
| 41 | 123241818 |
| 42 | 110731150 |
| 43 | 97307548 |
| 44 | 84019610 |
| 45 | 70493583 |
| 46 | 58467815 |
| 47 | 47395858 |
| 48 | 37756130 |
| 49 | 29430971 |
| 50 | 22613725 |
| 51 | 17277359 |
| 52 | 13029380 |
| 53 | 9656760 |
| 54 | 7277427 |
| 55 | 5348515 |
| 56 | 3918702 |
| 57 | 2911593 |
| 58 | 2155849 |
| 59 | 1659545 |
| 60 | 1220316 |
| 61 | 977314 |
| 62 | 789915 |
| 63 | 651484 |
| 64 | 532894 |
| 65 | 454101 |
| 66 | 387817 |
| 67 | 362678 |
| 68 | 356267 |
| 69 | 341987 |
| 70 | 305919 |
| 71 | 301722 |
| 72 | 273069 |
| 73 | 254269 |
| 74 | 236894 |
| 75 | 230284 |
| 76 | 207731 |
| 77 | 192670 |
| 78 | 155005 |
| 79 | 147265 |
| 80 | 144729 |
| 81 | 148739 |
| 82 | 132023 |
| 83 | 120266 |
| 84 | 107220 |
| 85 | 103838 |
| 86 | 103432 |
| 87 | 87471 |
| 88 | 88764 |
| 89 | 75777 |
| 90 | 67282 |
| 91 | 70530 |
| 92 | 66591 |
| 93 | 72294 |
| 94 | 66908 |
| 95 | 66570 |
| 96 | 66947 |
| 97 | 69066 |
| 98 | 63122 |
| 99 | 62349 |
| 100 | 71686 |
| 101 | 56445 |
| 102 | 54598 |
| 103 | 61734 |
| 104 | 54358 |
| 105 | 53423 |
| 106 | 51417 |
| 107 | 53848 |
| 108 | 52231 |
| 109 | 56100 |
| 110 | 47477 |
| 111 | 44084 |
| 112 | 45788 |
| 113 | 44938 |
| 114 | 41859 |
| 115 | 43022 |
| 116 | 39338 |
| 117 | 37570 |
| 118 | 39053 |
| 119 | 34331 |
| 120 | 29677 |
| 121 | 29614 |
| 122 | 26806 |
| 123 | 28374 |
| 124 | 29107 |
| 125 | 22088 |
| 126 | 20930 |
| 127 | 19299 |
| 128 | 16759 |
| 129 | 18671 |
| 130 | 16948 |
| 131 | 18268 |
| 132 | 12609 |
| 133 | 18447 |
| 134 | 12703 |
| 135 | 13415 |
| 136 | 13369 |
| 137 | 15640 |
| 138 | 10143 |
| 139 | 8249 |
| 140 | 12595 |
| 141 | 5854 |
| 142 | 5745 |
| 143 | 6763 |
| 144 | 4892 |
| 145 | 4104 |
| 146 | 3326 |
| 147 | 4558 |
| 148 | 3096 |
| 149 | 4633 |
| 150 | 3875 |
| 151 | 5150 |
| 152 | 3295 |
| 153 | 4180 |
| 154 | 3718 |
| 155 | 3572 |
| 156 | 2611 |
| 157 | 3705 |
| 158 | 4592 |
| 159 | 4397 |
| 160 | 4077 |
| 161 | 2085 |
| 162 | 3471 |
| 163 | 3160 |
| 164 | 3002 |
| 165 | 2742 |
| 166 | 2982 |
| 167 | 2944 |
| 168 | 2633 |
| 169 | 2295 |
| 170 | 2517 |
| 171 | 1366 |
| 172 | 2837 |
| 173 | 2032 |
| 174 | 2125 |
| 175 | 1759 |
| 176 | 1389 |
| 177 | 3016 |
| 178 | 3646 |
| 179 | 3603 |
| 180 | 1344 |
| 181 | 2432 |
| 182 | 2818 |
| 183 | 2465 |
| 184 | 1129 |
| 185 | 1939 |
| 186 | 1079 |
| 187 | 1213 |
| 188 | 863 |
| 189 | 720 |
| 190 | 352 |
| 191 | 148 |
| 192 | 680 |
| 193 | 1004 |
| 194 | 827 |
| 195 | 1068 |
| 196 | 1197 |
| 197 | 1742 |
| 198 | 377 |
| 199 | 1324 |

Figure S1. Coverage histogram for PacBio HiFi reads mapped to the PR1 genome. The x-axis shows the coverage and the y-axis shows the number of bases in the assembly at that coverage. The small bump at approximately X=18 represents reads from regions where the two haplotypes are divergent. Because the assembly only captures one haplotype, those regions show approximately half the coverage of the rest of the genome. The peak near X=0 represents regions in the PR1 assembly where minimap2 failed to align HiFi reads.

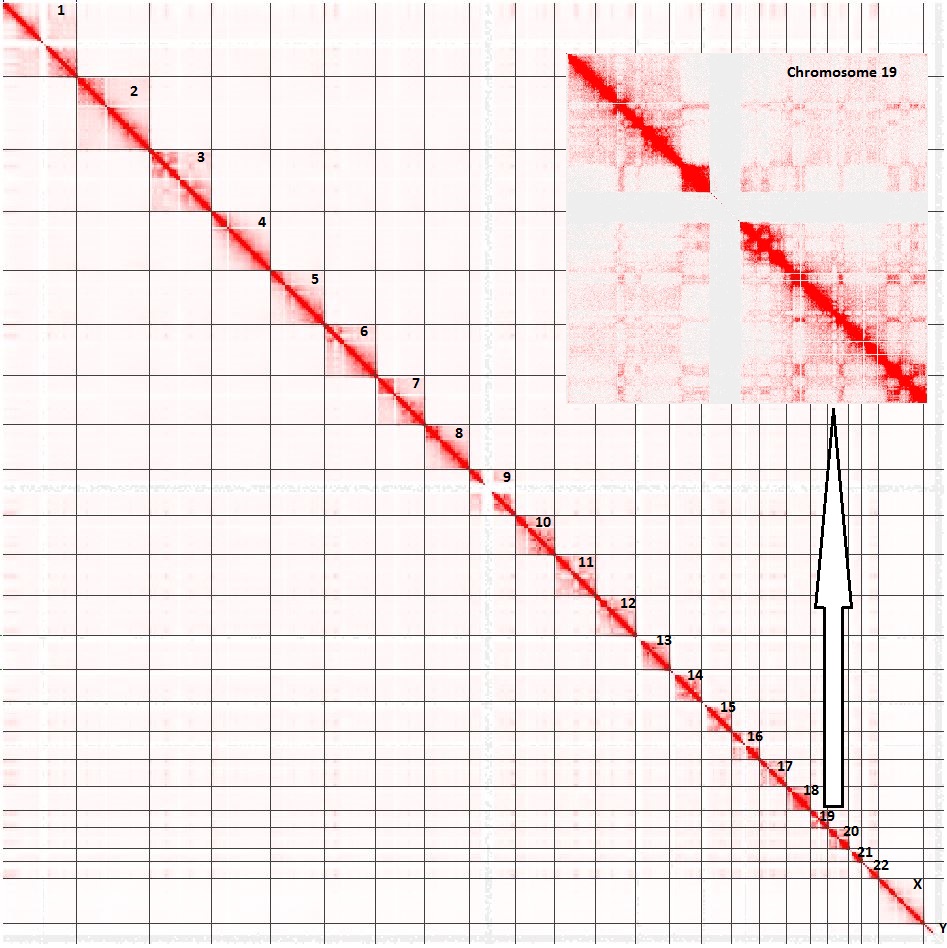

**Figure S2**. Hi-C contact map for the PR1 assembly. We used the Juicebox software to map 756,339,138 Illumina reads with average length of 123bp (after trimming) containing 92.7Gb of sequence to the PR1 genome. As shown here, most contacts cluster along the main diagonal, as expected for a correctly assembled genome. At the whole-genome scale, some of the smallest chromosomes are unclear, but a zoomed-in view, as shown in the inset for Chromosome 19, shows that the Hi-C data support the correctness of these chromosomes as well.
